# Genome-wide CRISPR screen reveals Wnt signaling defects regulate lipid accumulation in APOE4 oligodendrocytes

**DOI:** 10.1101/2025.10.16.682845

**Authors:** Leyla Anne Akay, Anna Bright, Carles A. Boix, Kate Louderback, Janine Medrano, Dingcheng Sun, Haonan Lin, Oisín King, Gwyneth M. Welch, Hiba Nawaid, Emre Agbas, Alan Jiang, Adele Bubnys, Ji-Xin Cheng, Joel Blanchard, Li-Huei Tsai

## Abstract

APOE4 is the largest genetic risk factor for late-onset Alzheimer’s disease, but the cellular mechanisms by which APOE variants influence risk of disease remain incompletely understood. We have previously found that APOE4 expression led to the intracellular accumulation of lipid droplets in oligodendrocytes, causing decreased myelination. However, the mechanisms by which APOE4 alters lipid metabolism are not fully understood. Here, we leveraged a genome-wide CRISPR screen and ATAC-sequencing in human induced pluripotent stem cell (iPSC)-derived oligodendrocytes to dissect APOE4’s lipid-associated mechanisms of action. Using these approaches, we identified decreased Wnt signaling, and overactive GSK3b activity, as regulators of lipid droplet accumulation in oligodendrocytes. Genetic and pharmacological inhibition of GSK3b reduced lipid droplets in APOE4 oligodendrocytes, and increased myelination in three-dimensional iPSC-derived brain organoids. Finally, we show that pharmacological inhibition of GSK3b reduces lipid droplets and improves myelination in APOE4;PS19 Tau transgenic mice. Together, our results provide a framework for understanding the mediation of APOE4-related changes to oligodendrocyte lipid metabolism and myelination.

## Introduction

Alzheimer’s disease (AD) is the most prevalent neurodegenerative disorder, affecting an estimated 7.2 million individuals in the United States^1^. With limited treatment options available, there exists an urgent need for the identification of novel therapeutic targets. Large scale genetic association studies have identified hundreds of genes contributing to risk of developing late-onset AD, with the strongest associations consistently found in the *APOE* gene locus^2^. *APOE* encodes Apolipoprotein E, a lipid transporter highly expressed in the brain. APOE exists across the human population largely as three variants: APOE3, the major allele; APOE2, a protective allele, and APOE4, a risk allele with regards to AD. One copy of APOE4 increases AD risk by an estimated 3-4 fold, whereas homozygosity increases risk by up to 12-fold^2^.

APOE4 has previously been implicated in a diverse suite of cellular processes, ranging from amyloid-beta aggregation, inflammation, neurofibrillary tau pathology and blood-brain-barrier breakdown^3^. However, an exact understanding of how APOE4 expression exerts its risk on AD remains elusive. We have previously performed single-nuclear RNA sequencing of the human post-mortem brain of individuals carrying the APOE4 allele, and found that APOE4 led to lipid and cholesterol dysregulation, most prominently in oligodendrocytes^4^. Oligodendrocytes are the myelinating cells of the central nervous system, responsible for providing electrical insulation to neuronal axons. Dysfunction of oligodendrocytes and breakdown of myelin is increasingly recognized not only as a feature of Alzheimer’s disease, but potentially a pathology-driving mechanism^5^. Therefore, understanding oligodendrocyte dysfunction and myelin maintenance within the contexts of aging and neurodegenerative disease is critical for the identification and development of new therapeutic opportunities.

## Results

### Genome-wide CRISPR screen reveals transcriptional programs underlying lipid accumulation in APOE4 oligodendrocytes

What are the molecular processes underlying the accumulation of lipid droplets in APOE4 oligodendrocytes? To gain a transcriptional handle on this disease-associated phenotype, we performed a genome-wide CRISPR knock-out screen in induced pluripotent stem cell (iPSC)-derived oligodendrocytes. We used a well-characterized iPSC line^6^, homozygous for the APOE4 allele, to generate oligodendrocytes, as previously described^7^. Upon differentiation, these iPSC-derived oligodendrocytes express canonical markers of mature oligodendrocytes, including transcription factor Olig2, and oligodendrocyte-specific surface antigen O4 (**Extended data figure 1A**).

We infected 90 million iPSC-derived APOE4 oligodendrocytes with an all-in-one guide RNA library containing Cas9 and targeting every protein-coding gene in the human genome, at a multiplicity of infection *circa* 0.3-0.5 (**Figure 1A**)^8^. The viral construct contains a puromycin resistance cassette, enabling selection of infected oligodendrocytes^8^. After antibiotic selection, oligodendrocytes were cultured for a further ten days, to allow enough time for the genomic lesion to affect protein turnover^8^. After these ten days, live oligodendrocytes were stained for lipid droplets using the fluorescent dye BODIPY, harvested and FAC-sorted into two populations: cells with the top 15% of BODIPY fluorescence (“high lipid burden”) and cells with the lowest 15% of BODIPY fluorescence (“low lipid burden”) (**Extended Data Figure 1B-F**). Genomic DNA was extracted from both populations, subjected to PCR amplification, and sequenced for guide RNAs (**Extended Data Figure 1G, H**). The enrichment of individual guide RNAs in both sorted populations were calculated relative to the original library pool, using the tool Apron (**Extended Data Figure 1I, J**).

**Figure 1:**
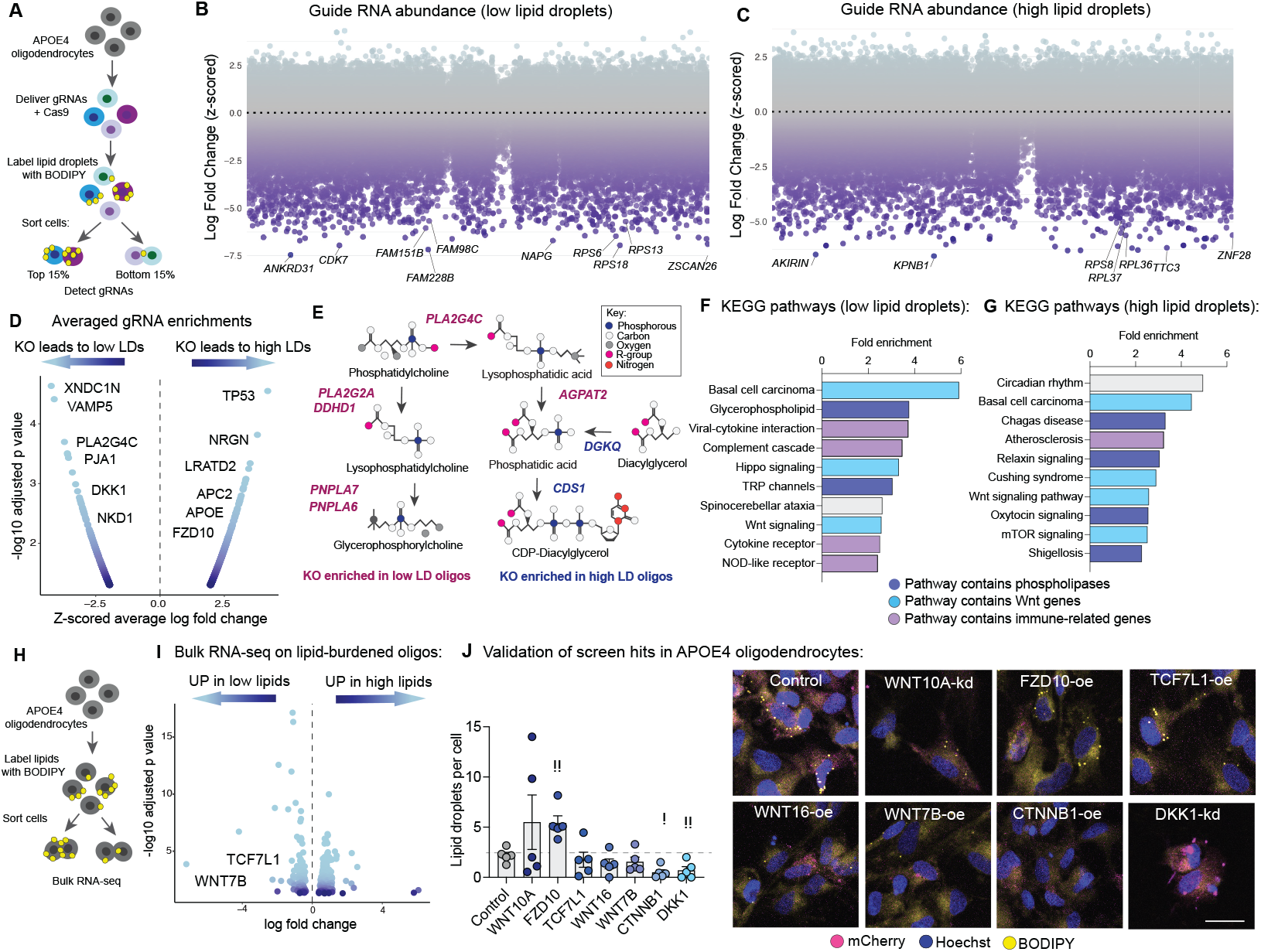
Genome-wide CRISPR knockout screen reveals regulators of lipid droplet burden in APOE4 oligodendrocytes. A, Schematic illustrating the experimental design. 90 million iPSC-derived oligodendrocytes were infected with a guide RNA library targeting every protein-coding gene in the genome at a multiplicity of infection circa 0.3-0.5. Ten days after infection, oligodendrocytes were stained for lipid droplets and FAC-sorted. Genomic DNA was harvested from both populations and used to calculate guide RNA enrichment. B, Manhattan plot illustrating the relative abundance of every gene knockout in oligodendrocytes with low lipid burden. Each dot represents the average abundance of all guide RNAs targeting an individual gene. Highly depleted genes are labeled. C, Manhattan plot illustrating the relative abundance of every gene knockout in oligodendrocytes with high lipid burden. Each dot represents the averaged abundance of all guide RNAs targeting an individual gene. Highly depleted genes are labeled. D, Volcano plot showing the z-scored enrichment of guide RNAs, averaged per gene. Genes highly enriched in the oligodendrocytes with low lipid droplets are shown on the left, and genes highly enriched in the oligodendrocytes with high lipid droplets are shown in the right. E, Schematic illustrating several of the screen hits whose enzymatic activity is related to phospholipid metabolism. F, Enriched KEGG pathways from significant hits in oligodendrocytes with low lipid droplets. G, Enriched KEGG pathways from significant hits in oligodendrocytes with high lipid droplets. Pathways are colored according to whether they contain genes related to phospholipases (purple), Wnt signaling (blue), or immune-related genes (pink). H, Schematic illustrating experimental design where APOE4 oligodendrocytes were labeled with BODIPY and FAC-sorted based on lipid content, and subjected to bulk RNA sequencing of both sorted populations. I, Volcano plot illustrating differentially expressed genes in oligodendrocytes with low (left) versus high (right) lipid burden. Labeled genes include Wnt-signaling related genes. J, Quantification and representative images of hit validation. APOE4 oligodendrocytes were infected with lentivirus to overexpress (oe) or knockdown (kd) Wnt related genes. Lipid droplets are labeled with BODIPY (yellow). Virus expression is shown with mCherry (pink). Nuclei labeled with Hoechst (blue). Scale bar represents 20 um. Each dot represents an individual experiment. This experiment was repeated several times.

As expected, both sorted populations showed highly significant depletion of guide RNAs targeting essential genes, including genes necessary for sustaining the Golgi apparatus, ribosome, and ubiquitin machinery. (**Figure 1B, C**). Among sorted populations, per-gene enrichments were calculated by averaging z-scored enrichments of individual guide RNAs (**Figure 1D**). In the “low lipid burden” population, significantly enriched genes included vesicular protein *VAMP5*, phospholipase *PLA2G4C*, two negative regulators of Wnt signaling, *DKK1* and *NKD1*, as well as 336 other genes (**Figure 1E, Supp. Table 1**). As expected, canonical lipid-droplet associated gene *PLIN3* was enriched, as were lipoprotein receptors *LRP2* and *VLDLR*. In the “high lipid burden” population, enriched genes included tumor suppressor *TP53*, lipid sensor *STARD10*, Wnt signaling related genes *APC2* and *FZD10*, and a further 401 additional genes (**Figure 1E**). Interestingly, *APOE* was also enriched, suggesting that loss of APOE-related lipoprotein function may provoke lipid droplet accumulation.

Several of our hits converged upon phospholipid metabolism (**Figure 1F**). Genes whose knockout were enriched in low lipid-burdened oligodendrocytes, i.e., genes likely to induce lipid accumulation, included phospholipases *PLA2G4C*, responsible for cleaving phosphatidylcholine to generate lysophosphatidic acid; *AGPAT2*, which converts lysophosphatidic acid to phosphatidic acid; *PNPLA6*, which cleaves phosphatidylcholine to generate glycerophosphorylcholine, and *PNPLA7*, which converts lysophosphatidylcholine to glycerophosphorylcholine. Notably, phospholipases have previously been reported to regulate lipid droplet formation^9^.

Conversely, the enzymes responsible for converting diacylglycerol to phosphatidic acid (*DGK1*), and phosphatidic acid to CDP-diacylglycerol (*CDS1*) were enriched in the high-lipid droplet population (**Figure 1F**). CDP-diacylglycerol is a substrate for the generation of phosphatidylinositols, which are key components of the plasma membrane and myelin sheath^10,11^. Together, these results suggests that enzymatic break-down of phosphatidylcholines and glycerophosphatdiylcholines by phospholipases may be a crucial step in lipid droplet formation in APOE4 oligodendrocytes, whereas the usage of diacylglycerols to generate phosphatidic acid, and potentially phosphatidylinositols, represents an alternative pathway.

To determine whether significantly enriched genes converged upon any known biological processes, we performed a gene set enrichment analysis. Among the “low lipid burden’ group, significantly enriched KEGG pathways included “basal cell carcinoma”, “glycerophospholipid metabolism,” and “viral-cytokine interaction” (**Figure 1G, Supp. Table 2**). Within the high lipid group, enriched pathways included “circadian rhythm,” “basal cell carcinoma,” and “Chagas disease.” (**Figure 1G, Supp. Table 3**). Several of the KEGG pathways were comprised of similar gene sets, including Wnt signaling genes (annotated as “basal cell carcinoma”, “Wnt signaling” and “Hippo signaling”), phospholipases, and immune-related genes. (**Figure 1G**). Crucially, although Wnt signaling was implicated in both high and low lipid populations, knock-out of the negative regulators of Wnt signaling (*DKK1, NKD1*) resulted in low lipid levels, whereas knock-out of canonical members of Wnt signaling (*FZD10, WNT16*) resulted in high lipid abundance.

To gain further insight into the transcriptional programs accompanying lipid burden in APOE4 oligodendrocytes, we next FAC-sorted naïve APOE4 oligodendrocytes (i.e., cells not subjected to any gene perturbations) based on lipid content (**Figure 1H**). We then performed bulk RNA-sequencing on the two sorted populations, comparing relative gene expression in low lipid-burdened versus high lipid-burdened oligodendrocytes. (**Extended Data Figures 1K-P**). APOE4 oligodendrocytes with increased lipid droplet burden upregulated a suite of lipid metabolic enzymes, including the cholesterol ester transferase protein *CETP*, ceramide synthase *CERS2*, and the enzyme *CDIPT* (CDP-diaclyglycerol transferase) (**Figure 1H, Supp. Table 4**). In contrast, oligodendrocytes with lower lipid burden had higher expression of diacylglycerol lipase alpha (*DAGLA*), an enzyme responsible for hydrolyzing diacylglycerols contained in lipid droplets. Notably, oligodendrocytes with lower lipid accumulation also had higher expression of Wnt-related genes *WNT7B* and *TCF7L1*, whereas oligodendrocytes with higher lipid accumulation upregulated *DAB2*, a negative regulator of Wnt signaling (**Figure 1H, Supp. Table 4**).

To validate our initial findings that Wnt signaling regulates lipid droplet accumulation in APOE4 oligodendrocytes, we next generated lentivirus to overexpress or knock-down (via shRNA) the Wnt signaling genes implicated in our CRISPR screen (*WNT10A, FZD10, WNT16, DKK1*), our RNA-sequencing (*WNT7B*), as well as the canonical master regulator of Wnt signaling, beta-catenin (*CTNNB1*) (**Figure 1I**). Ten days after infecting cultures of APOE4 oligodendrocytes, we labeled lipid droplets with BODIPY and used confocal microscopy to measure the number of lipid droplets per cell. We found that knock-down of *DKK1* significantly (p value 0.0155) reduced lipid droplet accumulation, as did overexpression of beta-catenin (*CTNNB1*) (p value 0.0031) (**Figure 1J**). Changes in gene expression were confirmed with RT-qPCR (**Extended Data Figure 1Q**). Taken together, these results provide evidence that Wnt signaling regulates lipid accumulation in APOE4 oligodendrocytes.

### ATAC-sequencing reveals transcriptional programs depressed in APOE4

To further understand how APOE4 perturbs the epigenome of oligodendrocytes, and to identify putative master regulators of APOE4’s effects on lipid accumulation, we next used ATAC-sequencing to compare transposase-accessible regions of the genome between iPSC-derived oligodendrocytes, isogenic except for APOE genotype (**Extended Data Fig. 2A, B**). Data quality was confirmed by calculating transcription start site enrichment scores (**Ext. Data Fig. 2C**) and fraction of reads within peak (**Ext. Data Fig. 2D**), which were both well within ENCODE standards.

**Figure 2:**
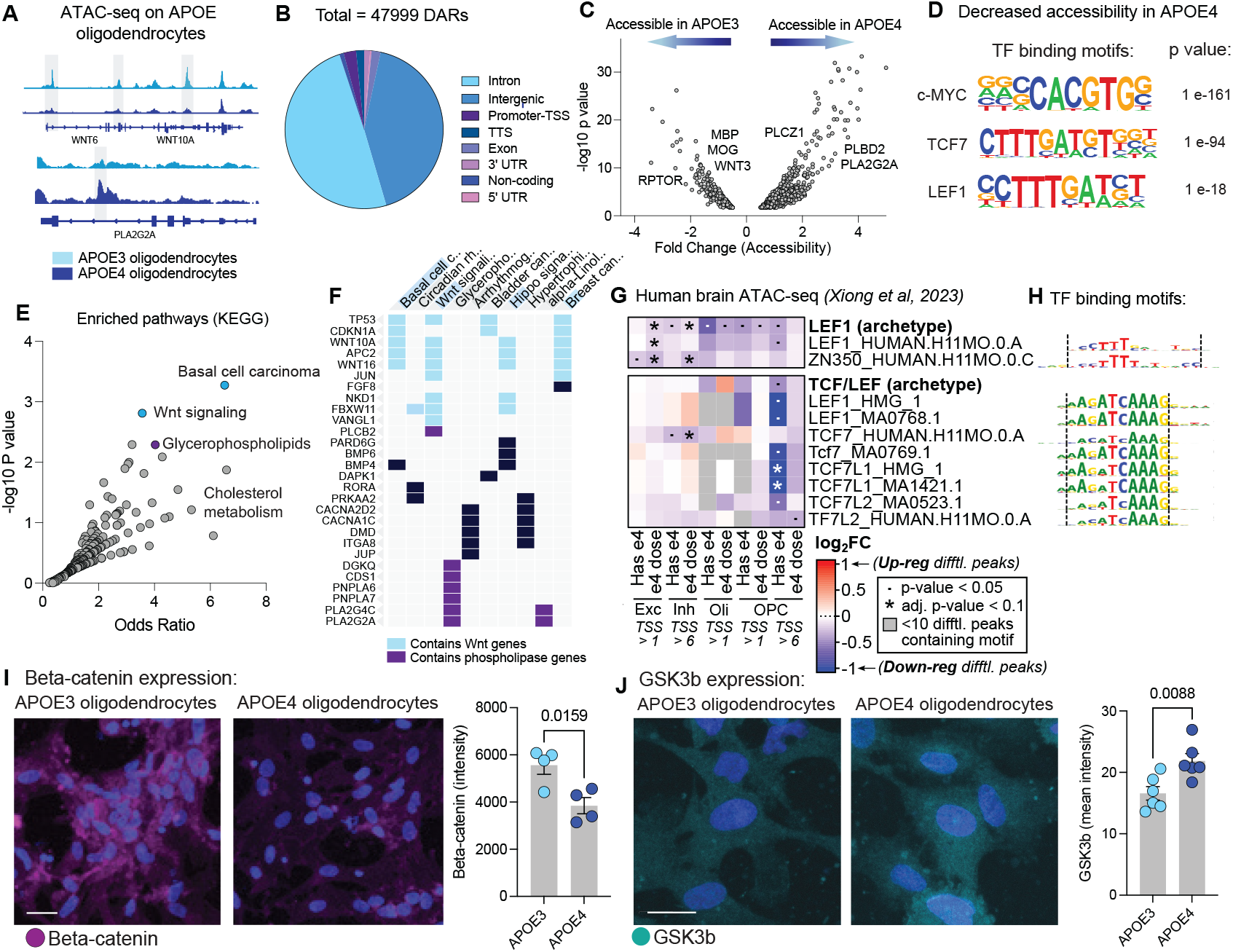
ATAC-sequencing reveals Wnt signaling is depressed in APOE4 oligodendrocytes. A, Pie chart illustrating the differentially accessible regions between APOE3 and APOE4 isogenic oligodendrocytes. B, Representative traces illustrating differentially accessible regions between APOE3 and APOE4 isogenic oligodendrocytes. C, Volcano plot showing the differentially accessible regions mapped onto protein-coding genes. D, Transcription factor binding motifs significantly enriched in regions of the genome less accessible in APOE4 oligodendrocytes. P value calculated using False Discovery Rate. E, Overlap between genes that are CRISPR screen hits and differentially accessible between APOE genotypes. This set of genes was queried for enriched KEGG pathways. F, Overlapping genes associated with enriched KEGG pathways. Pathways containing Wnt related genes are shown in blue. Pathways containing phospholipases are shown in purple. G, Transcription factor motif enrichment from ATAC-seq of the human post-mortem brain, comparing enrichment of Wnt motifs in differential peaks across individuals, stratified by e4 carrier status or e4 dosage. Cell types were evaluated at two TSS enrichment thresholds, only cell types with results with nominal p < 0.05 are shown. H, Logos for the enriched motifs, grouped into two archetypes. I, Representative images and quantification of immunoreactivity to beta-catenin (purple) n in cultures of isogenic APOE3 and APOE4 iPSC-derived oligodendrocytes. P value calculated using an unpaired, two-tailed student’s t-test. Each dot represents a separate culture. Scale bars represent 10 um. J, Representative images and quantification of immunoreactivity to GSK3b (cyan) in cultures of isogenic APOE3 and APOE4 iPSC-derived oligodendrocytes. P value calculated using an unpaired, two-tailed student’s t-test. Each dot represents a separate culture. Scale bars represent 20 um.

We identified a total of 47,999 differentially accessible regions between APOE3 and APOE4 oligodendrocytes, with the majority residing in introns and intergenic regions of the genome (**Figure 2A, Supp. Table 5**). After mapping the differentially accessible regions to genes, we identified a total of 9671 unique protein-coding genes with significant (false discovery rate-adjusted p value <0.05) differences in accessibility between APOE3 and APOE4 oligodendrocytes (**Figure 2B**). We observed decreased accessibility in genes related to Wnt signaling (*WNT3, TCF4, CTNNB1*, etc.), myelination (*MOG, CNP, MBP*), PI3K-Akt signaling (*RPTOR, IGF1, FGF1*), among others (**Figure 2C**). Genes with increased accessibility in APOE4 oligodendrocytes included phosphatidylcholine biosynthesis (*PCYT1A, PCYT1B, CHKA*), phospholipase metabolism (*PLBD2, PLCZ1, PLA2G2A*) and heparan sulfate biosynthesis (*HS3ST1, HS3ST4, HS6ST1*) among others (**Figure 2C**). Next, we used the tool *HOMER* to identify putative transcription factor binding domains enriched in differentially accessible regions (**Figure 2D, Supp. Table 6**). Motifs belonging to canonical Wnt transcription factors *TCF7, LEF1* and *c-MYC* were among those significantly enriched in regions less accessible in APOE4, suggesting a depression of canonical Wnt signaling in APOE4 (**Figure 2D**).

Lipid droplets can form in response to nutritional, inflammatory and even thermal cues^12^. Given that such distinct programs govern lipid droplet formation and breakdown across cell types, we wanted to narrow our focus onto APOE4-driven mechanisms. Therefore, we aimed to understand whether any of our CRISPR screen hits were also perturbed in APOE4 oligodendrocytes, relative to APOE3 oligodendrocytes. To test this, we compared differentially accessible regions between APOE3 and APOE4 with hits from our CRISPR screen, reasoning that the overlap would prioritize genes not only perturbed by APOE4, but also functionally related to lipid accumulation. We found 319 common genes (**Figure 2E, Supp. Table 7**), including members of the Wnt signaling family (*WNT10A, WNT, NKD1*) and several phospholipases (*PNPLA6, PNPLA7, PLA2G2A, PLA2G4C*). Pathway analysis of these overlapping genes again highlighted a significant enrichment of the terms “Basal cell carcinoma”, “Wnt signaling”, “Glycerophospholipids”, corresponding to canonical Wnt genes (*WNT10A, WNT16, NKD1*) and phospholipases (**Figure 2E, Figure 2F, Supp. Table 8**).

Given that Wnt signaling is known to play a role in neural development, we wanted to exclude the possibility that we were simply capturing differences in timing or development in our iPSC-derived models. Therefore, we next utilized a single-cell ATAC-sequencing dataset generated from human *post-mortem* prefrontal cortex, across 93 aged individuals^13^. Using this dataset, we called differentially accessible peaks in individuals carrying one or more copies of the APOE4 allele, compared to APOE3 homozygotes (**Figure 2G**). We calculated the enrichment of Wnt-related transcription factor binding sites within these differentially accessible peaks, and found a significant enrichment of Wnt signaling-related motifs in regions less accessible in APOE4 carriers, particularly in oligodendrocyte lineage cells and neurons (**Figure 2G, H**). This suggests that APOE4-related decreases in Wnt signaling can be detected in the context of the aging human brain.

Finally, to more directly test the hypothesis that Wnt signaling was perturbed in APOE4 oligodendrocytes, we performed immunohistochemistry against the protein beta-catenin, which is prevented from degradation in the presence of Wnt signaling^14^. We found a significant decrease in immunoreactivity to beta-catenin (p = 0.0159) in APOE4 oligodendrocytes compared to cultures of APOE3 oligodendrocytes, again suggesting that Wnt signaling was decreased in APOE4 oligodendrocytes (**Figure 2I**). In the absence of Wnt signaling, beta-catenin is phosphorylated by the kinase GSK3-beta, leading to its proteasomal degradation^14^. To determine whether decreased beta-catenin observed in APOE4 oligodendrocytes could be due to increased activity of GSK3-beta, we performed immunohistochemistry against GSK3-beta. We found a significant increase in immunoreactivity to GSK3-beta in APOE4 oligodendrocytes relative to APOE3, suggestive of increased GSK3-beta expression (**Figure 2J**).

Together, these results suggest that canonical Wnt/beta-catenin signaling is decreased in APOE4 oligodendrocytes, whereas GSK3-beta protein expression is elevated relative to APOE3.

### The beta-catenin / GSK3b axis regulates lipid droplet formation in oligodendrocytes

Our CRISPR screen highlighted Wnt signaling as a potential regulator of lipid droplet accumulation, and our ATAC-sequencing data showed that Wnt signaling was decreased in APOE4 oligodendrocytes. Therefore, we wanted to understand whether manipulation of Wnt signaling could be a tractable handle to modulate lipid droplet accumulation in oligodendrocytes. Wnt signaling is complex, with different Wnt ligands able to activate, repress, or interfere with downstream effectors in context-specific manners. Given our findings that beta-catenin was decreased and GSK3b had increased expression in APOE4 oligodendrocytes, and that beta-catenin overexpression led to highly significant reduction in lipid droplets, we decided to target the GSK3-beta / beta-catenin axis. GSK3b and beta-catenin are mutually repressive; GSK3b phosphorylates beta-catenin, tagging it for ubiquitination and proteasomal degradation^15^. Conversely, in the presence of beta-catenin-dependent Wnt signaling, GSK3b is rendered inactive by sequestration into multi-vesicular bodies^16^.

To directly test whether overactive GSK3-beta could drive lipid accumulation in the absence of APOE4, we first pharmacologically stimulated GSK3-beta activity in APOE3 oligodendrocytes using the small molecule Dif3^17^. We found that activating GSK3-beta in APOE3 oligodendrocytes led to robust lipid droplet accumulation (p value 0.0093) (**Figure 3A**). To test whether inhibiting GSK3-beta could reduce lipid accumulation in APOE4 cells, we used the drug CHIR99021 to inhibit GSK3-beta activity, and found that GSK3b inhibition significantly (p value 0.0014) reduced lipid droplet accumulation in APOE4 oligodendrocytes (**Figure 3B**). We additionally performed Stimulated Raman Scattering (SRS) microscopy, which enables the identification and quantification of lipid species in single cells^18^, on cultures of APOE4 oligodendrocytes with and without CHIR99021 treatment. Using this approach, we found that GSK3b inhibition led to a significant reduction in phospholipid and cholesterol content within individual lipid droplets (**Extended Data Figure 3A**).

**Figure 3:**
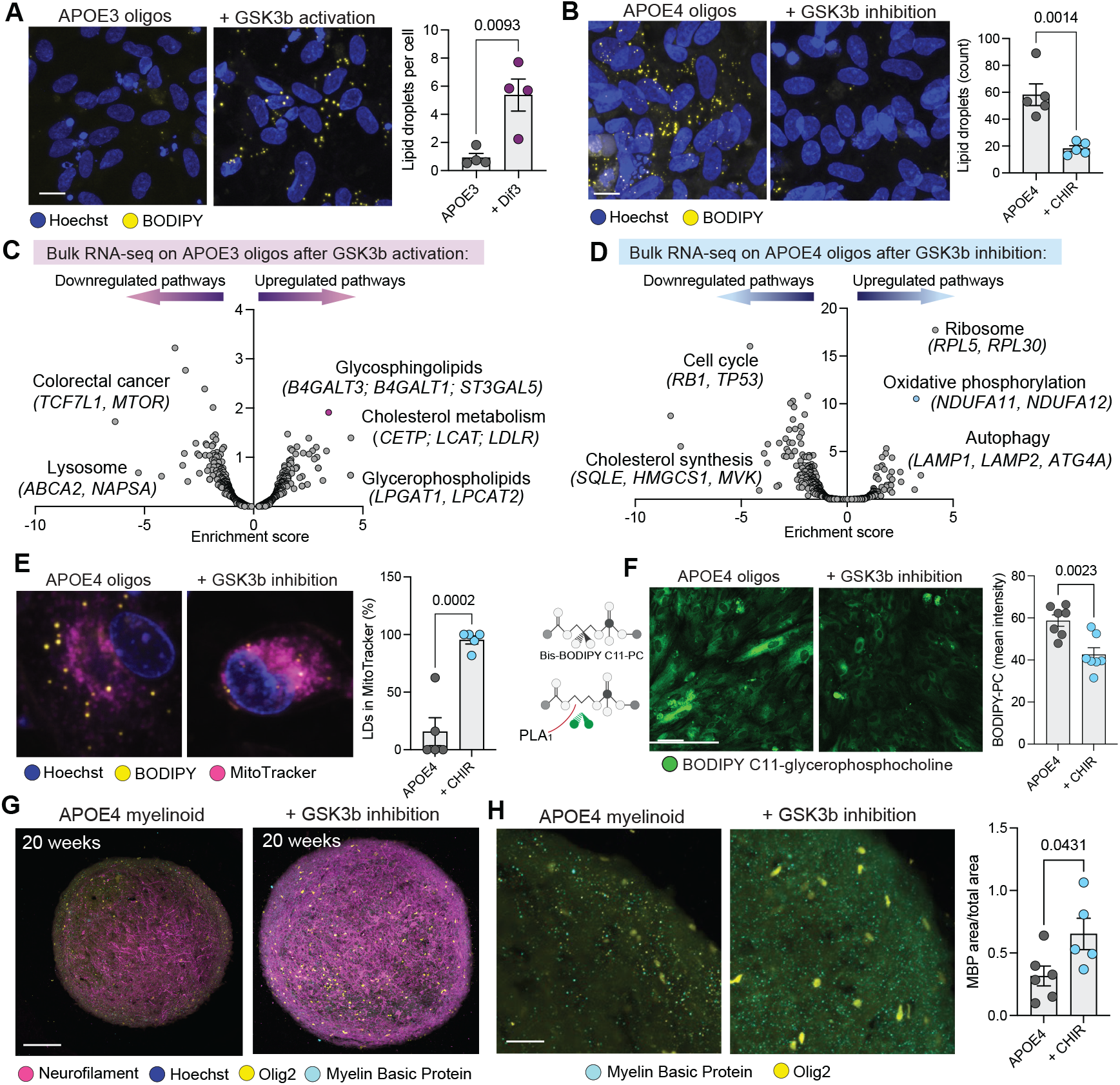
GSK3b activity regulates lipid droplet burden and myelination in oligodendrocytes. A, Treatment with GSK3b activator “Dif3” in APOE3 oligodendrocytes. Representative images and quantification of BODIPY (lipid droplet) staining in yellow; nuclei (Hoechst) in blue. Each dot represents an individual culture. P values calculated using two-tailed student’s t-test. Scale bar 10 microns. B, Treatment with GSK3b inhibitor CHIR99021 (1 uM) in APOE4 oligodendrocytes. Representative images and quantification of BODIPY (lipid droplet) staining in yellow; nuclei (Hoechst) in blue. Each dot represents an individual culture. P values calculated using two-tailed student’s t-test. Scale bar 10 microns. C, Volcano plot showing enriched KEGG pathways from bulk RNA-sequencing of APOE3 oligodendrocyte cultures treated with GSK3b activator “Dif3”. Relevant KEGG pathways are labeled. D, Volcano plot showing enriched KEGG pathways from bulk RNA-sequencing of APOE4 oligodendrocyte cultures treated with GSK3b inhibitor “CHIR99021”. Relevant KEGG pathways are labeled. E, Representative images of APOE4 oligodendrocytes treated with GSK3b inhibitor CHIR99021. Mitochondria are labeled with MitoTracker (magenta), lipid droplets are labeled with BODIPY (yellow) and nuclei with Hoechst (blue). Each dot represents and individual culture. P values calculated using two-tailed student’s t-test. F, Representative images and quantification of APOE4 oligodendrocyte cultures treated with GSK3b inhibitor CHIR99021. BODIPY C11-glycerophosphocholine is labeled in green fluorescence. Each dot represents an individual culture. P value calculated using two-tailed student’s t-test. G, Representative images of APOE4 myelinoids, showing control (left) and CHIR99021-treated (right). Axons are stained with neurofilament (magenta), myelin with Myelin Basic Protein (cyan), Olig2 (yellow) and nuclei with Hoechst (blue). Scale bar. H, Representative images and quantification of immunoreactivity to Myelin Basic Protein (cyan) in APOE4 control and CHIR99021-treated myelinoids. Each dot represents the average of three images from individual myelinoids. P value calculated using two-tailed student’s t-test.

To gain further insights into how GSK3-beta activity could be affecting lipid metabolism, we performed bulk RNA sequencing on cultures of APOE3 oligodendrocytes following GSK3-beta activation by Dif3 treatment. We observed broad upregulation of genes related to glycosphingolipid synthesis (*B4GALT3, B4GALT1, ST3GAL5*), glycerophospholipid metabolism (*PLA2G2E, LPCAT1, LPGAT1*) and cholesterol metabolism (*CETP, LCAT, LDLR*) (**Figure 3C, Supp. Table 9**). Notably, these results parallel the results from our initial CRISPR screen, which identified phospholipase enzymes as regulators of lipid droplet formation; *LPCAT1* is responsible for re-acylation of lysophospholipids. Both *CETP* and *LCAT* are critical regulators of lipid droplet formation, responsible for esterifying free cholesterol and packaging it into lipoprotein-like particles^19^.

Next, to determine how GSK3-beta inhibition could be decreasing lipid droplet burden in APOE4 oligodendrocytes, we performed bulk RNA-sequencing on APOE4 oligodendrocytes treated with CHIR99021. Among genes down-regulated after GSK3b inhibition, we noted a marked decrease of many genes involved in cholesterol biosynthesis: *HMGS1, MVK, MVD, SQLE, LSS, SREBF1* and *SREBF2*, genes we previously found were aberrantly upregulated in APOE4 oligodendrocytes^4^ (**Extended Data Figure 3B**). The most positively enriched KEGG pathways corresponded to “Ribosome” and “Oxidative phosphorylation”, consisting of many genes encoding the mitochondrial NADH dehydrogenase, cytochrome oxidase, and ATPase units (**Figure 3D, Supp. Table 10**), in agreement with Wnt signaling’s known effects on mitochondrial biogenesis and oxidative phosphorylation^20^. We additionally observed upregulation of *NPC1*, a transporter responsible for shuttling cholesterol to the lysosome, as well as upregulation of *LIPA*, a lysosomal lipase responsible for breaking down triglycerides and cholesterol esters. We also found broad upregulation of genes related to autophagy, including *LAMP1, LAMP2, ATG4A*, and *ATG14* (**Figure 3F**). Together, these results suggest that inhibition of GSK3-beta might reduce lipid droplet burden by promoting mitochondrial bioenergetic use of stored lipids by oxidative phosphorylation, or potentially lipophagy.

To experimentally test the hypothesis that GSK3b inhibition could promote mitochondrial usage of lipid droplets, we performed a pulse-chase experiment, in which we labeled mitochondria in APOE4 oligodendrocytes using the fluorescent mitochondrial dye MitoTracker, and lipid droplets using BODIPY. We then inhibited GSK3-beta in cultures by treatment with CHIR99021. Using confocal microscopy in live cultures, we then measured lipid droplet association with mitochondria (**Figure 3E**). Intriguingly, we found that GSK3b inhibition led to a significantly increased (p value 0.0002) association of lipid droplets with mitochondria (**Figure 3E**). To test whether GSK3b inhibition could similarly lead to lysosomal degradation of lipid droplets, we performed a similar experiment using the lysosome-specific fluorescent dye LysoTracker. GSK3b inhibition did not affect lipid droplet-lyososomal colocalization (**Extended Data Figure 3C**).

Given that we had originally identified dysregulated phospholipase activity in our CRISPR screen, and that our SRS data demonstrated a decrease in phospholipid content within lipid droplets, we next tested whether GSK3b inhibition could affect phospholipase activity (**Figure 3F**). We incubated APOE4 oligodendrocytes with BODIPY-C11-glycerophosphocholine. This molecule is a glycerophosphocholine, with a fluorescent BODIPY tag at the site of phospholipase-directed cleavage. When phospholipases cleave this substrate, the BODIPY is liberated, and green fluorescence can be detected. After treating APOE4 oligodendrocytes with CHIR99021, we detected a significant decrease in BODIPY-PC fluorescence, indicating reduced phospholipase activity (**Figure 3F**).

Oligodendrocytes are primarily responsible for generating and maintaining the myelin sheath. Therefore, to test whether restoring Wnt signaling via GSK3b inhibition could improve myelination, we turned to a three-dimensional iPSC-derived brain organoid model, containing myelinating oligodendrocytes (“myelinoids”)^21^. We generated these iPSC-derived myelinoids as previously described^21^. After 20 weeks in culture, myelinoids generated from APOE4 iPSCs had significantly reduced (p value 0.0012) myelination compared to those generated from isogenic APOE3 iPSC lines, as measured by immunoreactivity to myelin basic protein, suggesting that this model captures known APOE4-associated oligodendrocyte defects (**Extended Data Figure 3D**). After 18 weeks in culture, we began treating our APOE4 myelinoids with CHIR99021 (2 uM) (**Figure 3G**). After 2 weeks of CHIR99021 treatment, we performed immunohistochemistry on the myelinoids. We found that CHIR99021 treatment significantly (p value 0.0431) increased immunoreactivity to myelin basic protein (MBP) in APOE4 myelinoids (**Figure 3H**). We did not detect a significant difference in the number of Olig2-positive cells, indicating that this effect was likely not mediated through increases in oligodendrocyte proliferation, but potentially through increased myelin production from existing oligodendrocytes (**Extended Data Figure 3E**). We detected a significant (p value 0.0044) increase in immunoreactivity to neurofilament staining, indicative that GSK3b inhibition promoted axonal growth (**Extended Data Figure 3F**).

Together, this data suggests that GSK3b inhibition may reduce lipid droplet accumulation in APOE4 oligodendrocytes and promote myelination by suppressing dysregulated cholesterol synthesis, decreasing phospholipase activity, and engaging mitochondrial-lipid droplet interactions.

### Inhibition of GSK3b reduces lipid droplets and increases myelination in an APOE4;PS19 Tau mouse model

Our results suggest that dysregulated GSK3-beta could drive lipid accumulation in APOE4 iPSC-derived oligodendrocytes, and that inhibiting GSK3-beta can reduce lipid droplet burden, potentially by promoting mitochondrial usage of lipid droplets, suppression of phospholipase activity, as well as increasing myelination. To test whether inhibition of GSK3b would be sufficient to reduce lipid pathology in an *in vivo* context, we employed the Tau PS19 transgenic mouse line, crossed to humanized APOE3 or APOE4 mice^22^. The expression of APOE4 has previously been shown to worsen tau pathology, exacerbate inflammation and lead to oligodendrocyte dysfunction in transgenic mice^22,23^. In agreement with these previous findings, we detected significant (p value 0.0321) increases in phosphorylated tau pathology in APOE4;PS19 mice, compared to APOE3;PS19 mice at 6-7 months of age (**Extended Data Figure 4A**). We also detected a significant (p value 0.0273) decrease in immunoreactivity to myelin basic protein in the hippocampus of APOE4;PS19 mice as early as three months of age (**Figure 4A**). We did not detect any changes in Olig2-positive cell number in the hippocampus, suggesting that defects in myelination are not primarily driven by gross loss of oligodendrocytes (**Extended Data Figure 4B**).

**Figure 4.**
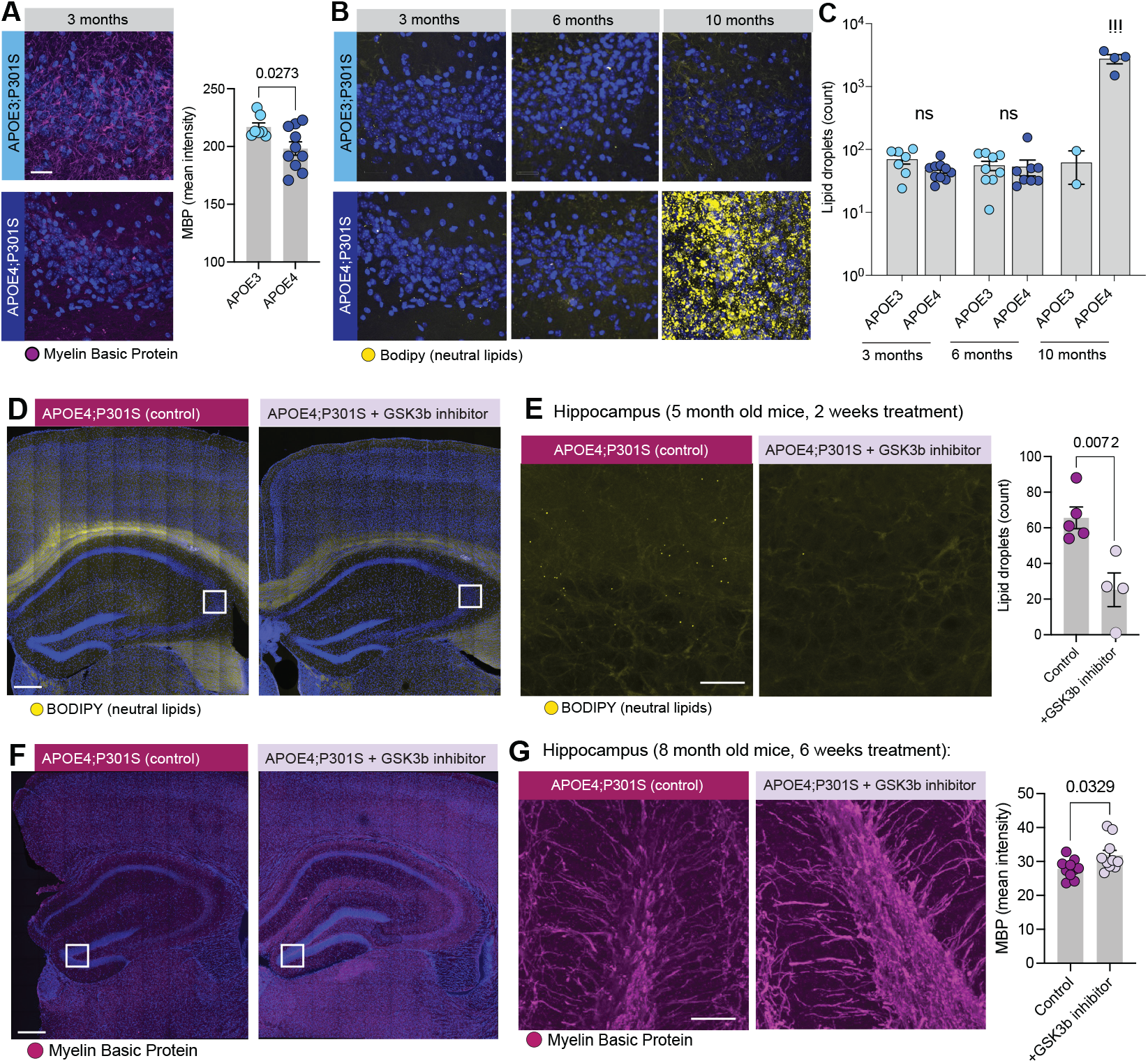
GSK3b inhibition reduces lipid droplets and increases myelination in an APOE4;Tau mouse model. A, Myelin basic protein immunohistochemistry (magenta) and nuclei with Hoechst (blue) in the hippocampus of APOE3;PS19 mice and APOE4;PS19 mice. Each dot represents an individual mouse. P value indicates two-tailed student’s t-test. B, Lipid droplet staining with BODIPY (yellow) and nuclei with Hoechst (blue) in the hippocampus of APOE3;PS19 and APOE4;PS19 mice at 3 months, 6 months and 10 months of age. C, Quantification of lipid droplet number in the hippocampus of APOE3;PS19 and APOE4;PS19 mice at 3, 6, and 10 months of age. Each dot represents an individual mouse. Statistical comparisons were performed by comparing genotypes at the same age using a two-tailed student’s t-test. D, Representative images showing lipid droplet staining (with BODIPY) in the hippocampus of control and treated APOE4;PS19 mice. E, Images and quantification of lipid droplets in the hippocampus of control (left) and treated (right) APOE4;PS19 mice. Each dot represents an individual mouse. P value was calculated using a two-tailed student’s t-test. F, Representative images showing immunoreactivity to myelin basic protein (magenta) in the hippocampus of control (left) and treated (right) APOE4;PS19 mice. G, Representative images and quantification of myelin basic protein immunoreactivity in control (left) and treated (right) APOE4;PS19 mice. Each dot represents and individual mouse. P value calculated using a two-tailed student’s t-test.

To determine whether APOE4;PS19 mice also harbored brain lipid accumulation, we stained for lipid droplets in hippocampal brain slices using BODIPY (**Figure 4B**). We found detectable lipid droplets throughout the hippocampus of both APOE3; and APOE4;PS19 mice, most notably in the CA3 region (**Figure 4B**). Although there were no measurable differences in the amount or localization of lipid droplets between APOE genotypes at three or six months of age, we found that APOE4;PS19 mice had a striking increase in lipid accumulation around ten months of age (**Figure 4C**).

To determine whether activation of Wnt signaling via reduction of GSK3b activity could reduce lipid droplets *in vivo*, we first treated a cohort of five-month old APOE4;PS19 mice with CHIR99021 at a dose of 5 mg/kg via intraperitoneal injection for two weeks (**Figure 5D**), before the onset of detectable tau pathology. Staining hippocampal brain sections for lipid droplets using BODIPY revealed a significant (p value 0.0072) reduction in lipid droplets in the hippocampus of CHIR-treated APOE4;PS19 mice, compared to vehicle-treated controls (**Figure 5E**). Next, to test whether restoring Wnt signaling over a longer time-course could increase myelination, we treated a cohort of eight-month old APOE4;PS19 mice with CHIR99021 for six weeks, and performed immunohistochemistry for myelin basic protein (**Figure 4F**). The longer-term treatment in older mice significantly increased hippocampal myelination (p value 0.0329), as measured by immunoreactivity to myelin basic protein (**Figure 4G**). We did not detect changes in Olig2-positive cell number (**Extended Data Fig. 4C**), immunoreactivity to GFAP (**Extended Data Fig. 4D**), number or morphology of microglia (**Extended Data Fig. 4E-F**), suggesting that the effects on myelination are primarily due to increased myelin production from existing oligodendrocytes. Given GSK3b’s well-documented role as a tau kinase, we also measured immunoreactivity to tau phosphorylated at Threonine 181 or Serine 396 residues, where GSK3b is known to phosphorylate tau. We did not detect a significant difference in immunoreactivity to tau phosphorylation at either residue (**Extended Data Fig. 4G, H**).

## Discussion

Here, we use a genome-wide genetic screen to identify regulators of lipid droplet accumulation in APOE4 oligodendrocytes, and find an unexpected role of the canonical Wnt signaling pathway. We find that Wnt signaling is depressed in APOE4 oligodendrocytes, both in iPSC-derived models and in the aged human post-mortem brain, and that deregulated Wnt / GSK3b kinase activity is involved in the development of lipid accumulation. Importantly, restoration of Wnt signaling via GSK3b inhibition reduced lipid droplet accumulation and increased myelination in both human iPSC-based models, as well as transgenic mouse models.

Aberrant lipid accumulation has been reported in a range of APOE4-related contexts, from transgenic mouse models^24^, to iPSC-derived microglia^25,26^, astrocytes^27,28^ and oligodendrocytes^4^. Lipid accumulation in these cell types has been linked to a variety of phenotypes, including cellular growth defects^28^, dysregulated neuronal activity^25^, defective oligodendrocyte maturation^29^ and decreased myelination^4^. Importantly, APOE4-associated glial lipid accumulation is also closely associated with neuronal tau pathology. Conditioned media from iPSC-derived microglia with high lipid burden was shown to provoke tau hyperphosphorylation in iPSC-derived neurons^26^. Promoting microglial cholesterol efflux by overexpression of lipid transporter ABCA1 or dietary treatment of LXR agonist was also found to reduce tau pathology in APOE4;PS19 mice^24^. Therefore, identifying pathways regulating lipid burden in APOE4 glia may advance the development of therapeutics targeting neuronal tau pathology.

Previous research has utilized both genome-wide and targeted CRISPR screens to identify regulators of lipid burden in APOE4 microglia, highlighting the contributions of triglyceride metabolism, lysosomal function and mTOR signaling^26,30^. Intriguingly, our genome-wide CRISPR screen in APOE4 oligodendrocytes recovered an enrichment of phospholipase enzymes. As these enzymes are primarily involved in cleaving phosphatidylcholines, the majority of which reside within cellular membranes, it is tempting to speculate that lipid droplets in APOE4 oligodendrocytes may be generated by aberrant cleavage of membrane-associated phospholipids, rather than uptake of exogenous fatty acids and subsequent conversion to triacylglycerols.

We also find that a major upstream pathway driving lipid accumulation in APOE4 oligodendrocytes is disrupted Wnt/beta-catenin signaling. Previous research has highlighted an intriguing relationship between APOE variants and Wnt signaling, with exposure to ApoE4 protein demonstrated to reduce Wnt/beta-catenin signaling in cell lines^31^, and variants in *LRP6* reported to increase Alzheimer’s disease risk in an APOE-genotype dependent manner^32^. Of note, the rare protective APOE variant “Christchurch” was recently shown to increase Wnt / beta-catenin signaling in human iPSC-derived brain organoids^33^. The potential convergence of rare protective variants and common risk variants in APOE onto changes in Wnt signaling certainly warrants further investigation.

The kinase GSK3b is a major inhibitor of Wnt signaling, and here we find that inhibition of GSK3b activity significantly lessens lipid droplet burden in APOE4 oligodendrocytes, both in human cells and transgenic mouse models. Given GSK3b’s established role as a major tau phosphorylating kinase, our work draws an interesting potential parallel between oligodendrocyte lipid accumulation and neuronal tau pathology. Recent work has demonstrated that reduced lithium bioavailability contributes to Alzheimer’s disease onset and progression in aged individuals, through activation of GSK3b^34^. Notably, disrupting Wnt signaling by provoking lithium deficiency in a mouse model of AD not only exacerbated tau pathology, but also had profound consequences on oligodendrocyte function, including demyelination^34^. Together, these findings implicate GSK3b as a shared pathological connection between glial lipid accumulation and neuronal tau pathology, highlighting its relevance to Alzheimer’s disease etiology.

## Acknowledgements

We thank David Root and members of the Broad Institute’s Genetic Perturbation Platform for assistance with CRISPR screening. We thank Stuart Levine, Duanduan Ma and members of the BioMicroCenter for assistance with RNA and ATAC sequencing and analysis. We thank Mat Victor for assistance with iPSC culture. We thank Ying Zhou, Ute Geigenmuller and Erica McNamara for administrative support and assistance with mouse colony management. This work was partially supported by NIH grants RF1AG075901, UH3NS115064 to L-H T, the Robert and Renee E Belfer Family Foundation, the Manthripragada McManama Alzheimer’s Fund, CAMA Seed Fund, the Picower Institute for Learning and Memory, J. Crayton Pruitt Foundation, Joseph P. DiSabato and Nancy Sakamoto.

## Declaration of Interests

A patent application has been filed by LAA and LHT on the results of the CRISPR screen. LHT is a member of the scientific advisory board of 4M Therapeutics and TAC Therapeutics.

## Materials

Key Resource Table

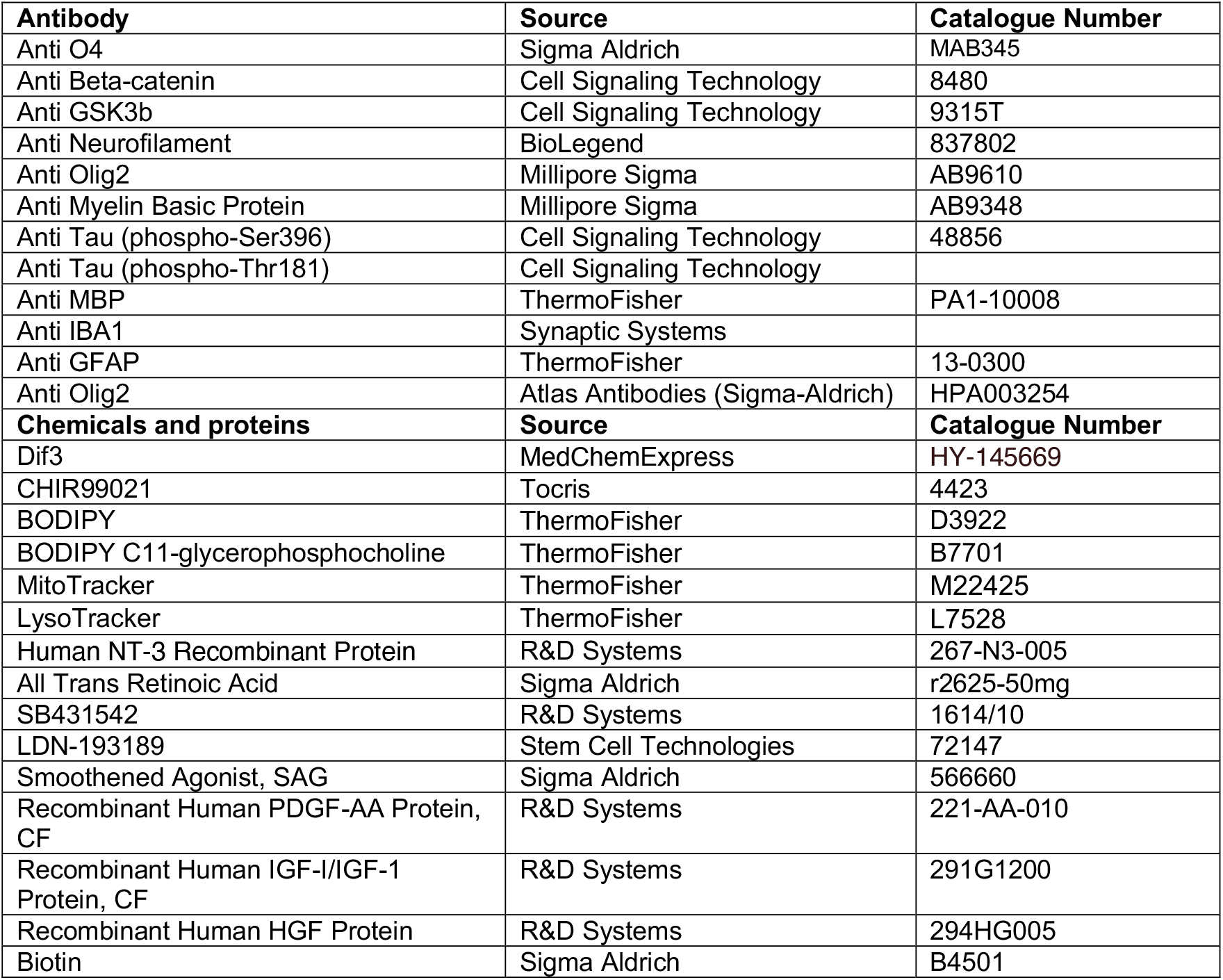
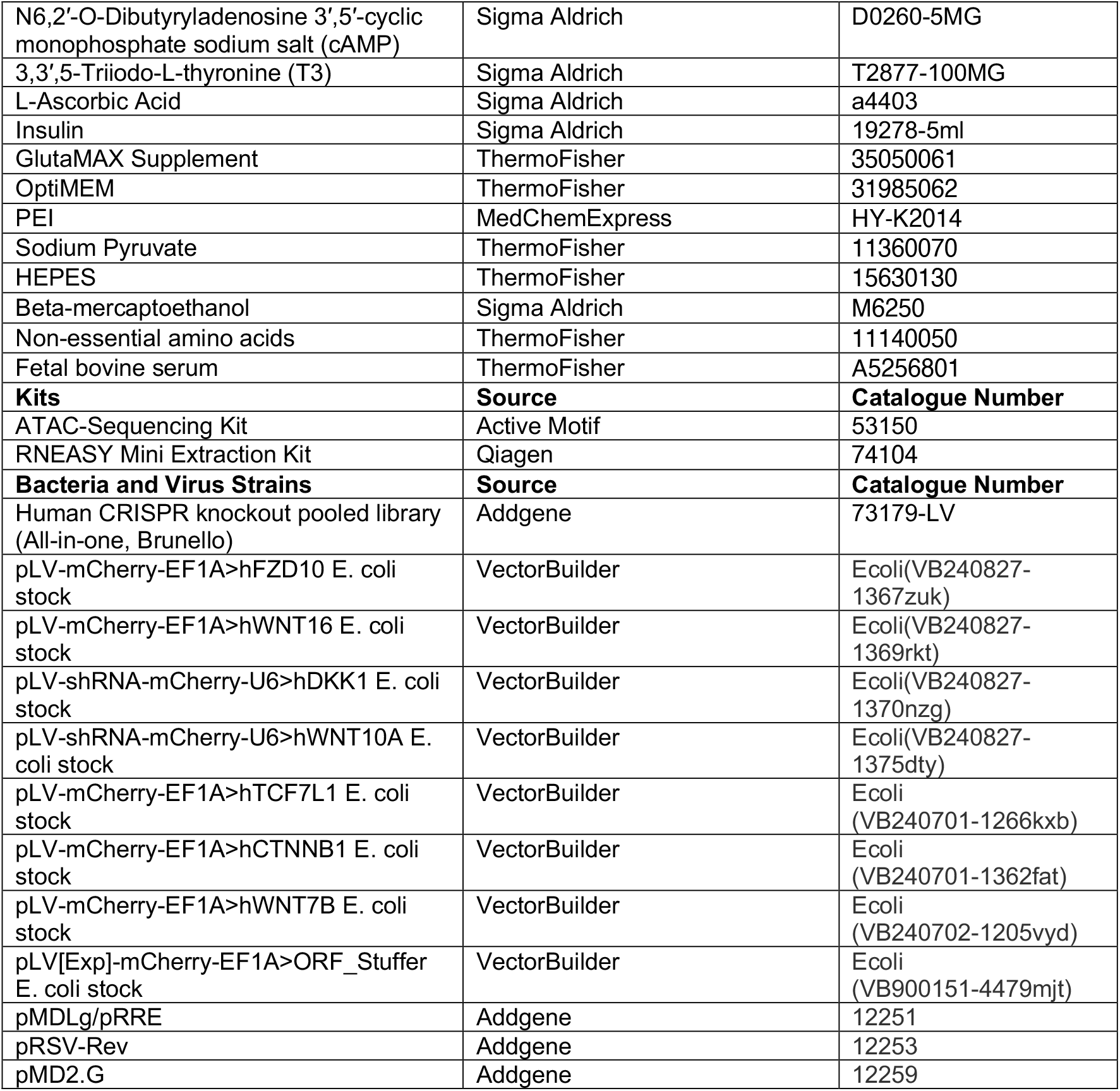

## Methods

### iPSC-derived oligodendrocyte differentiation

Oligodendrocytes were generated according to previously established protocols^7^. Briefly, colonies from isogenic APOE3 and APOE4 induced pluripotent stem cells (iPSC)^6^ were plated into 6-well plates, and cultured until colonies reached approximately 250 um in diameter. Cells were then cultured in “Neural Induction” media, mTSER containing 10 μM SB431542, 250 nM LDN193189 and 100 nM RA. Media was changed every day. On day 8, the media was changed to “N2 media”, or DMEM/F12 media containing non-essential amino acids, GlutaMAX, N2 supplement, 2-mercaptoethanol, Penicillin-Streptomycin, with 100 nM RA and 1 μM SAG added each day. On day 12, aggregated cells were mechanically lifted and re-plated into ultra-low-attachment 10 cm dishes. Cell aggregates were then cultured in N2B27 media. On day 30, the aggregates were plated into 6-well plates coated with Poly-L-orinthine and laminin, to enrich for oligodendrocyte precursors. Oligodendrocyte precursors were then cultured in N2/B27 media containing PDGFaa (10 ng/ml), IGF-1 (10 ng/ml), HGF (5 ng/ml), NT3 (10 ng/ml), T3 (60 ng/ml), Biotin (100 ng/ml), cAMP (1 μM) and Insulin (25 μg/ml). On day 75, cells were then cultured in N2/B27 media containing HEPES (10 mM), T3 (60 ng/mL), Biotin (100 ng/mL), cAMP (1 uM), Insulin (25 ug/mL), and Ascorbic acid (20 ug/mL). After day 75, oligodendrocyte cell fate was validated by immunohistochemistry for canonical markers of lineage (Olig2) and maturity (O4).

### Genome-wide CRISPR knockout screen

90 million APOE4 iPSC-oligodendrocytes were plated into 150 mm dishes, at a density of approximately 9 million cells per dish. Cells were infected with the Brunello Human CRISPR Knockout pooled library, containing Cas9, at a multiplicity of infection between 0.3 and 0.5. After 36 hours, cells were treated with Puromycin to enable selection for infected oligodendrocytes. After a subsequent ten days in culture, cells were treated with BODIPY to label lipid droplets. 24 hours after BODIPY treatment, media was changed to remove excess BODIPY, and cells were lifted into FACS buffer and subjected to FAC-sorting based on green fluorescent signal. After gating for exclusion of dead cells, doublets and debris, single cells with the top 15% and the lowest 15% of green fluorescent signal were sorted separately and collected. Genomic DNA from each population was extracted using the NucleoSpin Blood XL Maxi kit, and guide RNAs were subjected to PCR amplification and sequencing. Guide RNA enrichment was calculated relative to the original library pool of DNA, using the Broad Institute’s tool “Apron”. Genes were deemed significantly “enriched” or “depleted” if the z-scored average of their corresponding guide RNAs’ log fold change had an adjusted p-value lower than 0.05.

### RNA-sequencing on sorted populations

APOE4 iPSC-oligodendrocytes were treated with BODIPY overnight. The next morning, media was changed and cells were collected and subjected to FAC-sorting based on BODIPY signal. The top 15% and the lowest 15% of green fluorescent signal were separately sorted, and cells were pelleted. The cell sorting was repeated in triplicate. RNA was extracted from all populations, using Qiagen’s RNeasy Kit, according to manufacturer’s instructions. Bulk RNA-sequencing was then performed on sorted populations of APOE4 iPSC-oligodendrocytes, using a 50 + 50 bases pair-end run with 8 + 8 nucleotide indexes, on a Singular 50PE run with 100m reads, at MIT’s BioMicroCenter core facility.

### Lentivirus generation and gene manipulation

HEK293T cells were plated in 10 cm tissue culture dishes, in DMEM media containing Glutamax, 10% Fetal Bovine Serum, 1% Non-essential amino acids, 1% Sodium Pyruvate, 1% HEPES buffer, 1% Pen/Strep solution, 0.1% beta-mercaptoethanol. The next day, 10 ug of each transfer vector, 5 ug pMDLg, 2.5 ug pRSV-Rev, 2.5 ug pMD2.G, 600 uL of OptiMEM, and 48 uL of PEI were added to Eppendorf tubes, inverted gently and incubated at room temperature for 20 minutes. This transfection solution was then added to HEK293T cells drop-wise. Media was changed the following day. Two days later, lentivirus-containing media was collected from dishes, and centrifuged at 1200 xg for 5 minutes at 4°C. The supernatant was poured into a Millex 0.45 um PES filter cup into ultracentrifuge tubes, and spun down in a Beckman ultracentrifuge for 2 hours at 25,000 rpm using a SW32Ti rotor. At the end of the spin, media was aspirated and the viral pellet was resuspended in 100 uL of phosphate-buffered saline. The lentiviral pellet was aliquoted and stored in −80°C. Viral aliquots were only used once, and never thawed and re-frozen. APOE4 oligodendrocytes were plated into 6-well plates at a density of 250,000 cells per well. Cells were treated with 10 uL of virus per well. 10 days after transfection, viral expression was assessed by mCherry expression. Cells were then treated with BODIPY, and re-plated into either 96-well plates for confocal imaging, or 6-well plates for RNA extraction.

### qPCR analysis of viral gene manipulation

Following RNA extraction and purification, Bio-Rad’s iScript™ Reverse Transcription Supermix (#1708840) was used to create cDNA, per manufacturer’s instructions. cDNA was quantified using a NanoDrop Spectrophotometer ND-1000. The qPCR was carried out according to the protocol provided by Bio-Rad for the SsoFast™ EvaGreen® Supermix (#1725201), and the Bio-Rad CFX96 System was used to perform the cycling per the optimized conditions suggested in the protocol. The number of cycles was increased from 40 to 50 to ensure sufficient amplification of all samples. A final concentration of 500 nM was used per primer per replicate, and 50 ng of cDNA was used per replicate. Once the samples were run in triplicate, the average Cq was found. The expression fold change for each sample was found using the following formula, 2^-((test experimental Cq - house experimental Cq) - (test control Cq - house control Cq)), where test refers to the primer used being that of the gene in question and house being the primers of the reference gene-in this case GAPDH. Experimental means the cDNA used is from cells infected with a virus to overexpress a certain gene, and control means the cDNA used is from the stuffer control, where the cells were infected with a virus with a random sequence instead of one of the target genes.

### ATAC-sequencing on APOE oligodendrocytes

ATAC-sequencing was performed using the ActiveMotif ATAC-sequencing kit, according to manufacturer’s instructions with minor modifications. Briefly, isogenic APOE3 and APOE4 oligodendrocytes were harvested and pelleted into LoBind DNA Eppendorf tubes, with 250,000 cells per replicate, and two replicates per genotype. Media was aspirated, and cells were centrifuged in 100 uL of ice-cold phosphate-buffered saline for five minutes at 500 rcf at 4C. Supernatant was discarded and cells were resuspended in 100 uL of ice-cold ATAC lysis buffer, and centrifuged for 10 minutes at 500 rcf at 4C. 50 uL of transposomes and tagmentation buffer mastermix was then added to each sample, and cells were resuspended. The tagmentation reaction was incubated at 37 C for 30 minutes, shaking at 800 rpm. 250 uL DNA purification binding buffer with sodium acetate was added to each sample, and DNA was purified and eluted. Tagmented DNA was then PCR amplified and subjected to SPRI size selection. Libraries were quantified and paired-end Illumina sequencing was performed by MIT’s BioMicroCenter. ATAC-sequencing data was first aligned to the human genome hg38 using the tool BowTie, and peaks were called using Macs2. The universal peakset was sorted and merged, and a peak count matrix was created using BedTools. The tool HOMER was used to assign peaks to nearest genes, using the “annotatePeaks.pl” command. Differentially accessible regions were calculated using DESeq2. Transcription factor binding motifs were identified using HOMER, with the “findMotifsGenome.pl” command, using parameters of size 200, and using the peak universe raw counts count matrix as the background.

### Analysis of ATAC-sequencing on post-mortem human brain

In order to find peaks in snATAC-seq of post-mortem samples of the prefrontal cortex of aged individuals, we aggregated snATAC-seq fragments from 93 individuals into fragment files for each of seven major cell groups (astrocytes, microglia, oligodendrocytes, OPCs, vascular cells, and excitatory and inhibitory neurons), keeping cells passing QC at each of two different TSS enrichment thresholds (TSS > 1 and TSS > 6), as previously reported^13^. We then removed unmappable and problematic regions using the umap kmer=50 annotation^35^ and the hg38 ENCODE blacklist (ENCFF356LFX)^36^. We called peaks with MACS3^37^ using “--call-summits --nomodel --extsize 147 -g hs -B -q 0.05” and counted the reads in peaks at the individual level using bedmap^38^ with --fraction-map 0.2. We used edgeR^39^ to perform differential peak calling in two conditions, presence of the APOE e4 allele and dosage (0/1/2) of the e4 allele in an individual, controlling for age, post-mortem interval, and sex. For differential peaks, we normalized using between-lane normalization in EDASeq^40^, filtered for expressed peaks, and estimated dispersion using estimateGLMCommonDisp in edgeR. We called differential peaks as all peaks with adjusted p-value < 0.05. To find motif instances in the hg38 genome, we applied MOODS^41^ with parameters “--p-value 4^(-motif_length) --lo-bg 2.977e-01 2.023e-01 2.023e-01 2.977e-01” for a set of 1,690 motifs comprised of JASPAR 2018 core non-redundant vertebrate motifs^42^, HOCOMOCO v1156^43^, and SELEX motifs by Jolma et al^44^. We partitioned the motifs into 286 previously identified motif archetypes^38^. To annotate peaks with motif instances, we intersected peaks with motifs using bedmap with --fraction-map 0.2. To test for motif enrichment in differential peaks, we first separated out up-regulated and down-regulated peaks for a given covariate, cell type group, and TSS QC threshold. We used a hypergeometric test to assess enrichment by comparing the number of peaks with a given motif in the differential set, the number of peaks with a motif in the overall peak set, the number of peaks in the differential set, and the total number of peaks in that cell type and QC threshold. We tested enrichment both for motifs and motif archetype, counted when seeing any of its constituent motifs, and adjusted p-values using the Benjamini-Hochberg correction within a peakset across all motifs and archetypes. We plot the enrichment in either up or down-regulated peaks depending on which is more significant or has a greater effect size (in that order), and flipped the sign of the log2FC for an enrichment in down-regulated peaks.

### Immunohistochemistry on APOE oligodendrocytes

Oligodendrocytes were plated into clear-bottomed 96 well plates at a density of 10,000 cells per well. Cells were fixed with 4% paraformaldehyde for twenty minutes at 4°C. Cells were washed with ice-cold phosphate buffered saline (PBS) twice for ten minutes, and then blocked in buffer (PBS containing 0.03% Triton X and 1% Bovine Serum Albumin) for two hours at room temperature, gently shaking. Cells were incubated in primary antibody solution (Rabbit anti-GSK3-b, 1:500, or Rabbit anti-beta-catenin, 1:500) overnight at 4°C, gently shaking. Cells were washed with ice-cold PBS three times for ten minutes, and then incubated in secondary antibody solution (Donkey anti-Rabbit Alexa Fluor 595, 1:1000; Hoechst, 1:10000) for two hours at room temperature, gently shaking. Cells were then washed three times with ice-cold PBS and imaged on a Zeiss LSM 900 confocal.

### Drug treatment on APOE oligodendrocytes

APOE3 and APOE4 oligodendrocytes were plated into separate clear-bottomed 96 well plates, at a density of 10,000 cells per well. APOE3 oligodendrocytes were treated either with Dif3 (10 uM) or vehicle control (DMSO). APOE4 oligodendrocytes were treated with the GSK3b inhibitor CHIR99021 (1 uM) or vehicle control (DMSO). Media containing drug was changed every other day. After one week of treatment, cells were treated with BODIPY to label lipid droplets. The following day, media was changed and cells were fixed with 4% paraformaldehyde, for 20 minutes at 4C. Cells were subsequently washed with ice-cold PBS twice for ten minutes, and then imaged with a Zeiss LSM 900 confocal microscopy.

### Organelle imaging in APOE4 oligodendrocytes

APOE4 oligodendrocytes were plated into a clear-bottomed 96-well plate at a density of 10,000 cells per well, and allowed to recover for three days. Cells were then treated with MitoTracker and BODIPY overnight. The following morning, media was changed, and cells were imaged using a live confocal microscopy system on the Zeiss LSM 900, with a heated stage supplying continuous CO2 at 5%. All conditions were imaged using the same laser power and microscopy settings.

### Organoid generation and immunohistochemistry

Oligodendrocyte-containing organoids (“myelinoids”) were generated as previously described, from isogenic iPSC lines (Parental KOLF2.1J, homozygous for APOE3, and an isogenic line, CRISPR edited to be APOE4 homozygous). Myelinoids were grown in culture for 18 weeks, at which point some APOE4 myelinoids began treatment with CHIR99021 (2 uM in media). Myelinoids were fixed in 4% PFA for 3 hours and submerged sequentially in 10%, 20%, and 30% sucrose overnight at 4 degrees. After full submersion in 30% sucrose, spheroids were embedded in the OCT compound and serially sectioned at 40 μm thickness using RWD Minux FS800 cryostat. Myelinoid sections were permeabilized in PBS with 0.1 % Triton X-100 (PBST) for 15 min and then blocked with 5% donkey serum in PBS (blocking buffer) for 3 hours at room temperature. Sections were then incubated with primary antibodies in blocking buffer at 4°C overnight. Following primary antibody incubation and three 10 minute washes, sections were incubated with secondary antibodies at room temperature for 2 hours, counterstained with Hoechst, washed three times for 10 minutes, and mounted with Fluoromount aqueous mounting medium.

### Visible hyperspectral SRS imaging

Visible hSRS system^45^ utilized a dual-output laser (Insight X3, Spectra Physics) where the output wavelengths of 906 nm and 1045 nm were used to obtain 453 nm and 523 nm through second harmonic generation. Hyperspectral imaging was achieved using the spectral focusing method^45^. We acquired the hyperspectral imaging data for the cells in the CH wavenumber region with 523 nm Stokes beam at 30 mW and 453 nm Pump beam at 20 mW. Furthermore, we applied Self-Supervised Elimination of Non-Independent Noise (SPEND) denoising to the hyperspectral data. SPEND is a self-supervised deep learning denoising framework for removing non-independent noises in hyperspectral images. Details can be found in^46^. Then, we used the LASSO method for the spectral unmixing of the hyperspectral stack into chemical maps of lipid, protein, cholesterol, and RNA^47^, using a least square fitting problem with L1-norm regularization with the parameter λ used to control each chemical component’s sparsity level.

### APOE4;PS19 mouse experiments

All experiments were done in accordance with MIT’s Institute for Animal Care and Usage Committee. All mice were housed in a 12-hour light/dark cycle room, with access to food and water *ad libitum*. All mice were housed with littermates. APOE4;PS19 male mice (five months old) were injected intraperitoneally with sterile CHIR99021 (5 mg/kg, 100 uL) or 10% DMSO twice a week for two weeks. Separately, APOE4;PS19 female mice (eight months old) were injected intraperitoneally with sterile CHIR99021 (5 mg/kg, 100 uL), or 10% DMSO twice a week for six weeks. One day after the final injection, mice were anesthetized with gaseous isoflurane and transcardially perfused with ice-cold phosphate-buffered saline (PBS). Brains were dissected out and drop-fixed in 4% paraformaldehyde for 16 hours. Brains were subsequently re-hydrated in PBS, and sliced on a Leica vibratome at a thickness of 40 um. Floating sections containing hippocampus were collected for immunohistochemistry in 24-well plates. For lipid droplet staining, to preserve lipid structures as much as possible, sections were not permeabilized with blocking buffer. Instead, brain sections were directly incubated in PBS containing BODIPY (1:1000) and Hoechst (1:10,000) for two hours at room temperature, and then washed three times with ice-cold PBS. Sections were mounted on Fisher Scientific Superfrost Plus glass slides, with VWR coverslips (22×30 mm No.1). Sections were then imaged on a Zeiss LSM 900 confocal. Lipid droplets were quantified using the “Analyze Particles” feature of ImageJ FIJI. Mean intensity of immunohistochemical stains was analyzed using the “Measure” feature of ImageJ FIJI. Microglial morphology was analyzed using the “filament” feature of IMARIS software. All experimenters were blinded to genotype and treatment condition of the mice.

**Extended Data Figure 1.**
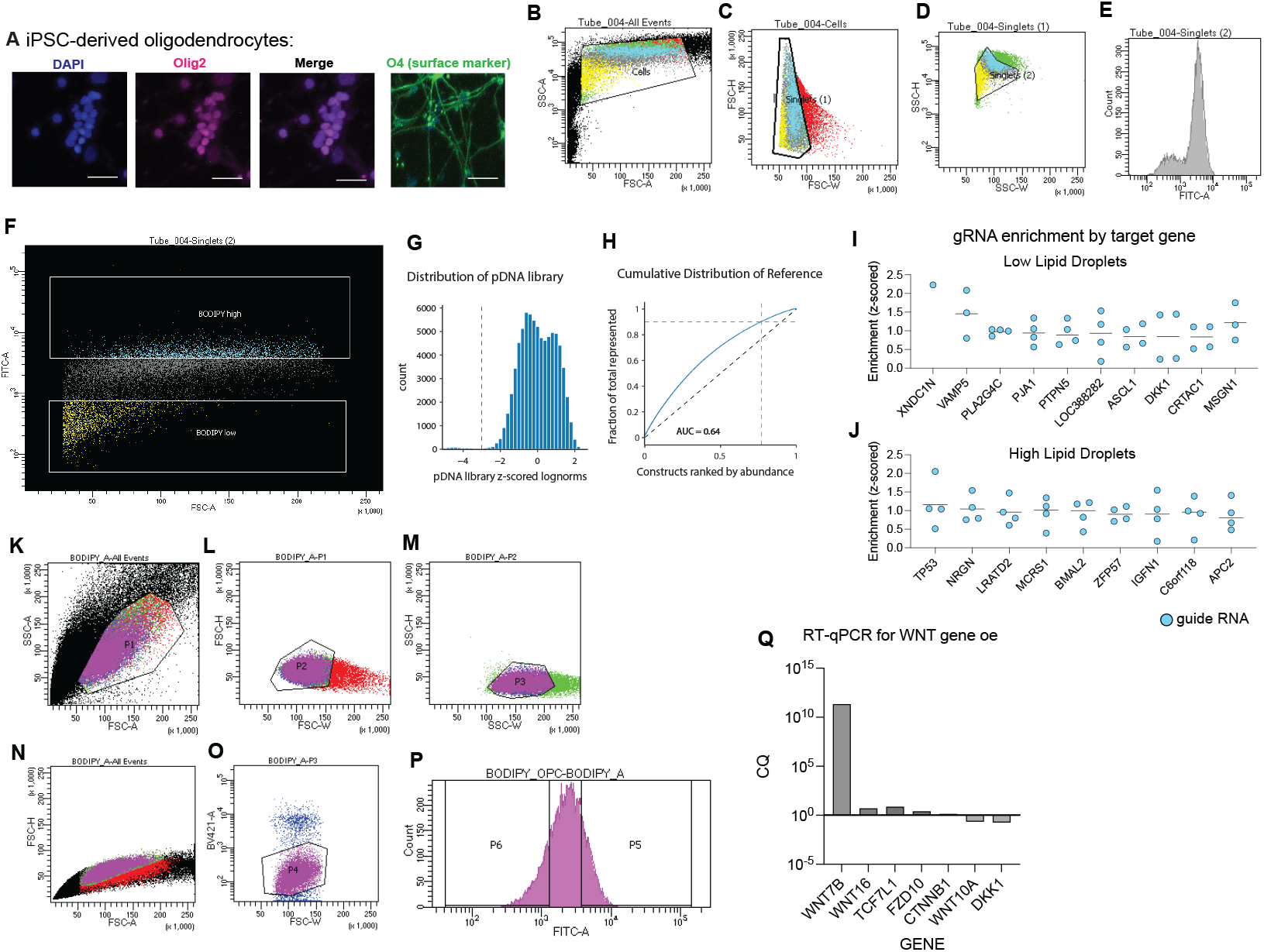
A, Representative images showing immunoreactivity to oligodendrocyte lineage markers in iPSC-derived oligodendrocytes. B, FACS plots showing gating for live cells. C, Gating for debris. D, Gating for singlets. E, Distribution of BODIPY (FITC-A) signal in cells. F, Distribution of cells with high(top) and low (bottom) BODIPY signal. G, Distribution of plasmid DNA in reference library. H, Cumulative distribution of reference library. I, J, Guide RNA enrichment per target gene for the top ten most enriched genes in cells with low lipid droplets (top) and high lipid droplets (bottom). Each dot represents an individual guide RNA. K, FACS plots showing gating for cells. L, Gating for Debris. M. Gating for singlets. N, Gating for. P, FACS plot showing distribution of BODIPY signal in sorted cells. Q, CQ values for RT-qPCR for gene expression of manipulated genes.

**Extended Data Figure 2.**
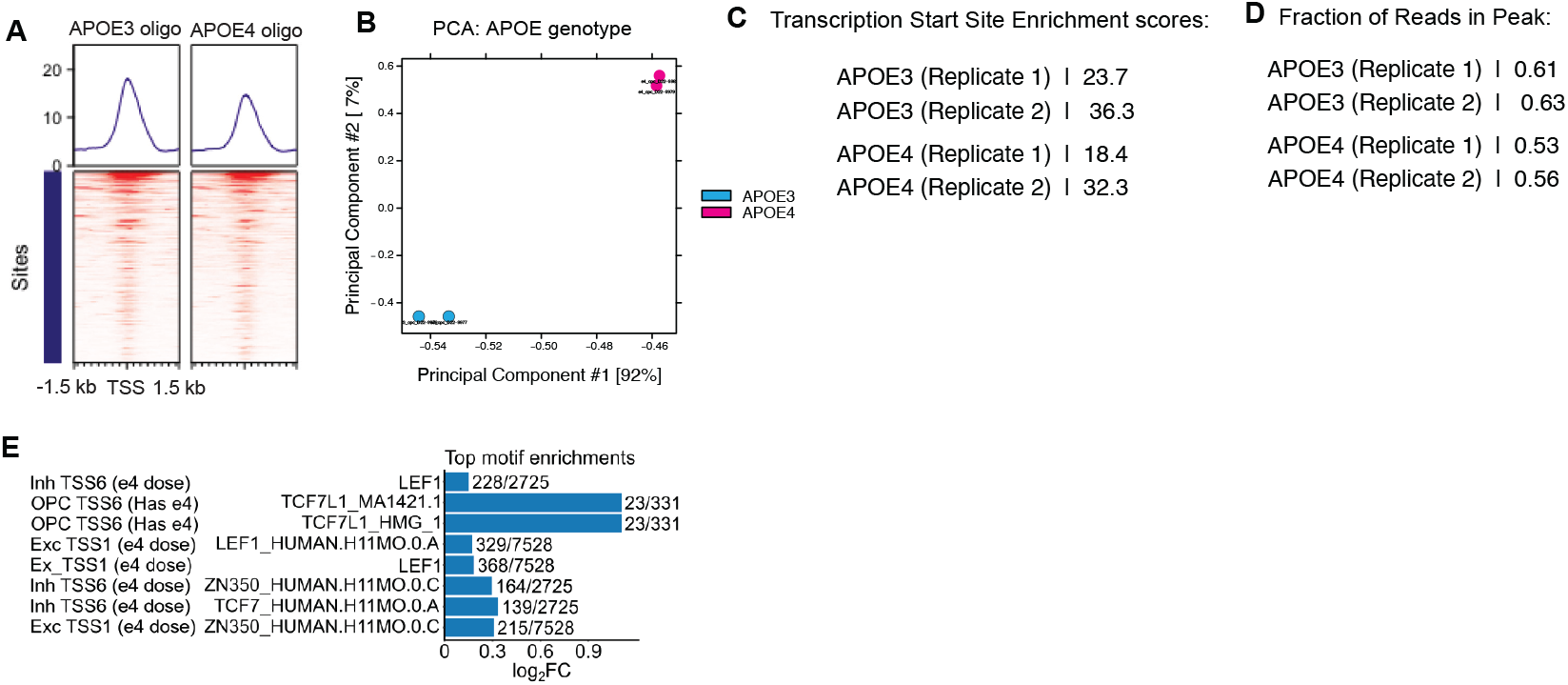
A, Genome-wide accessibility in isogenic APOE3 (left) and APOE4 (right) iPSC-derived oligodendrocytes. B, PCA plot showing ATAC-sequencing. Replicates cluster by APOE genotype. C, Transcription start site enrichment scores of each replicate used for ATAC-sequencing. D, Fraction of reads in peaks calculated for each replicate used for ATAC-sequencing. E, Log2 fold change for significant enrichments of Wnt motifs in cell type-specific differential peaks for e4 dosage and carrier status. Bars are labeled with the number of differential peaks with a motif over the total number of differential peaks.

**Extended Data Figure 3.**
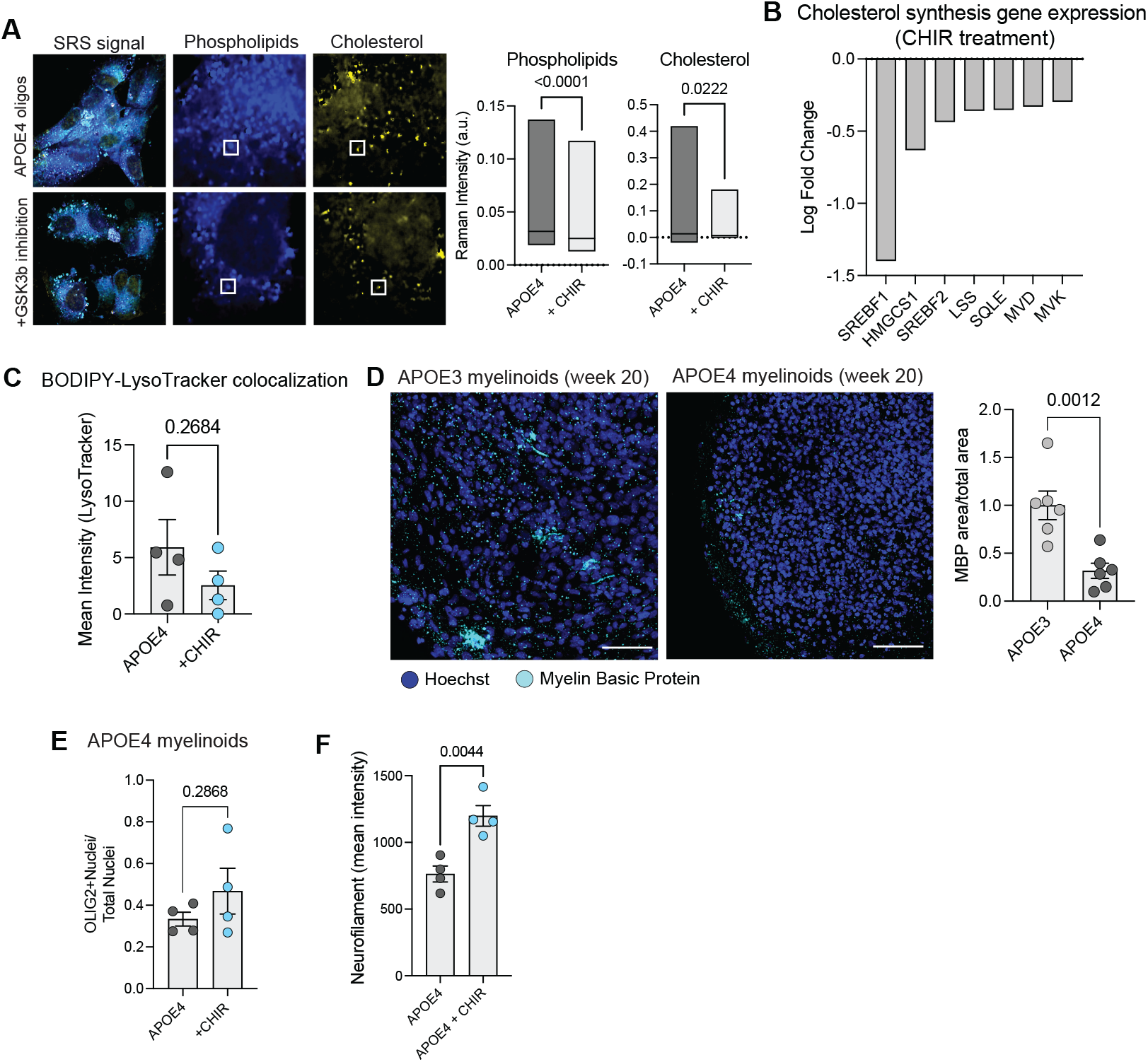
A, Stimulated Raman Scattering on cultures of APOE4 iPSC-derived oligodendrocytes, showing control (top) and CHIR99021-treated (bottom). SRS signal is shown for phospholipids (middle row) and cholesterol (right row). Raman intensity was calculated per lipid droplet. B, Gene expression (log fold change) of canonical cholesterol synthesis enzymes genes, in APOE4 oligodendrocytes with and without CHIR99021 treatment. C, Quantification of BODIPY (lipid droplet) and LysoTracker co-localization in APOE4 oligodendrocytes with and without CHIR99021 treatment. P value calculated using two-tailed student’s t-test. D, Myelin basic protein (cyan) staining in APOE3 myelinoids and APOE4 myelinoids after 20 weeks in culture. Scale bar represents 20 um. Each dot represents an individual myelinoid, P value calculated using a two-tailed student’s t-test. E, Number of Olig2-positive nuclei in APOE4 myelinoids with and without CHIR99021 treatment. Each dot represents an individual myelinoid. P value calculated using two-tailed student’s t-test. F, Neurofilament staining (mean intensity of immunoreactivity) in APOE4 myelinoids with and without CHIR99021 treatment. Each dot represents an individual myelinoid. P value calculated using two-tailed student’s t-test

**Extended Data Figure 4.**
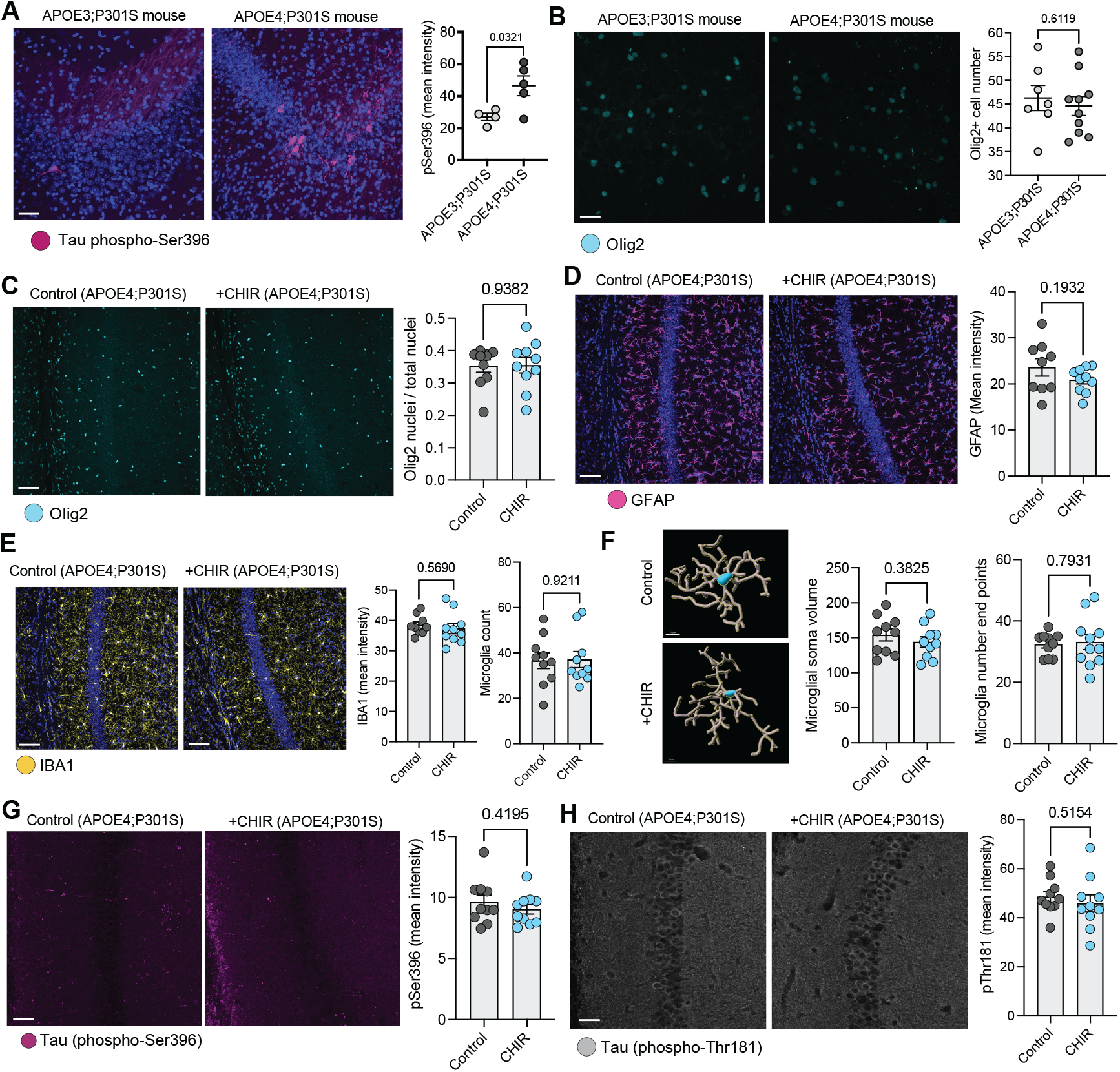
A, Representative images and quantification of phospho-tau Ser396 immunoreactivity in the hippocampus of APOE3;PS19 (left) and APOE4;PS19 (right) mice. Each dot represents an individual mouse. P value calculated using two-tailed student’s t-test. B, Representative images and quantification of number of Olig2-positive cells in the hippocampus of APOE3;PS19 (left) and APOE4;PS19 (right) mice. Each dot represents an individual mouse. P value calculated using two-tailed student’s t-test. C, Representative images and quantification of number of Olig2-positive cells in the hippocampus of APOE4;PS19 mice treated with control (left) or CHIR99021 (right). Each dot represents an individual mouse. P value calculated using two-tailed student’s t-test. D, Representative images and quantification of number of immunoreactivity to GFAP the hippocampus of APOE4;PS19 mice treated with control (left) or CHIR99021 (right). Each dot represents an individual mouse. P value calculated using two-tailed student’s t-test. E, Representative images and quantification of immunoreactivity to IBA1 and number of microglia in the hippocampus of APOE4;PS19 mice treated with control (left) or CHIR99021 (right). Each dot represents an individual mouse. P value calculated using two-tailed student’s t-test. F, Representative images showing 3-D reconstruction of microglial morphology in control (top) and treated (bottom) mice. Quantifications of microglial soma volume and number of processes. Each dot represents the averaged values of microglia detected per individual mouse. P value calculated using two-tailed student’s t-test. G, Representative images and quantification of phospho-tau Ser396 immunoreactivity in the hippocampus of APOE4;PS19 mice treated with control (left) or CHIR99021 (right). Each dot represents an individual mouse. P value calculated using two-tailed student’s t-test. H, Representative images and quantification of phospho-tau Thr181 immunoreactivity in the hippocampus of APOE4;PS19 mice treated with control (left) or CHIR99021 (right). Each dot represents an individual mouse. P value calculated using two-tailed student’s t-test.

